# Clodronate liposomes untangle the role of hemocytes in *Apis mellifera* response to temperature variation and microbial infection

**DOI:** 10.1101/2025.09.23.678096

**Authors:** Michael Oeth, Deepak Kumar, Michael Goblirsch, Mohamed Alburaki, John Adamczyk, Shahid Karim

## Abstract

**Introduction:** The cellular immune response, mediated by hemocytes, is a fundamental component of honey bee (*Apis mellifera*) health. However, the specific contribution of hemocyte subtypes to resilience against combined stressors remains poorly characterized. This study investigated the effect of temperature and bacterial challenge on hemocyte abundance. We employed clodronate liposomes (CLD) for the first time in honey bees as a novel tool for the selective depletion of hemocytes to investigate this gap.

**Methods:** Five-day-old (nurses) and fifteen-day-old (foragers) honey bees were treated with CLD, control liposomes, PBS, or left untreated, then exposed at either 32°C or 22°C and challenged with the gram-negative bacterium, *Escherichia coli,* or the gram-positive bacterium, *Staphylococcus aureus*. Survival, hemolymph volume, total hemocyte counts, and differential hemocyte counts were monitored over seven days from the start of exposure.

**Results:** The CLD application demonstrated significant reductions in granulocyte and prohemocyte populations, indicating the highest vulnerability. A temperature drop to 22°C buffered the negative impact on survival of CLD-induced immunosuppression. While bacterial challenges universally reduced hemocyte counts, we found an age-dependent difference where nurses maintained significantly higher baseline total hemocyte counts than foragers. Furthermore, temperature did not affect overall total hemocyte counts in 5-day-old nurse bees, but in 15-day-old foragers, it significantly modulated the hemocyte response to bacterial infection.

**Conclusion:** Our findings show that hemocyte function is non-uniform, with specific subtypes being essential for overall resilience. The results highlight a previously underappreciated role for temperature as a key modulator of immune capacity, particularly in immunocompromised bees. The age-related differences in hemocyte abundance suggest a life-history trade-off that may prompt the increased vulnerability of honey bees as they age. This work establishes CLD as a powerful tool for insect immunology and sets a precedent for using precise immune manipulation to study host-pathogen-environment interactions.

## INTRODUCTION

Honey bees (*Apis mellifera*) are essential pollinators, supporting both global ecosystems and agricultural productivity (Aizen et al., 2009). However, their populations are increasingly threatened by pathogens, parasites, and environmental stressors (Abban et al., 2024; Alburaki et al., 2024; Dennis & Kemp, 2016; Potts et al., 2010; Watanabe, 2008). Healthy, long-lived bees rely on their immune system, which protects against pathogenic microbes. Unlike vertebrates, invertebrates like honey bees lack an adaptive immune system and depend entirely on innate immunity, supplemented by immune priming mechanisms (Chan et al., 2009; Evans & Spivak, 2010; Gätschenberger et al., 2013; Hernández López et al., 2014; Leponiemi et al., 2021, 2023; Simone et al., 2009). Bees also contribute to social immunity by removing infected (Spivak and Reuter, 2001) or dead bees (Visscher, 1983) from the hive and collecting resin (propolis) with antimicrobial properties (Simone et al., 2009; Cremer et al., 2007). These defenses are crucial for maintaining colony health, especially under environmental pressures such as pesticide exposure, pathogen spread, habitat loss, and temperature extremes.

The innate immune system of honey bees consists of cellular and humoral components that act together to combat infections. Cellular immunity is mediated by hemocytes, which circulate in the hemolymph and perform key functions such as phagocytosis, encapsulation, nodulation, and wound healing (Evans et al., 2006; Gábor et al., 2020; Hystad et al., 2017; Richardson et al., 2018). Hemocytes detect pathogens through interactions between pathogen-associated molecular patterns (PAMPs) and pattern recognition receptors (PRRs), initiating immune responses such as phagocytosis and melanization (Evans et al., 2006; GarcÃ-a-Lara et al., 2005; Negri et al., 2013, 2014, 2016). Granulocytes, the primary phagocytic cells, engulf and digest microbial invaders, while plasmatocytes mediate encapsulation of larger pathogens or pathogen aggregates. Prohemocytes function as precursors, giving rise mainly to granulocytes and plasmatocytes to maintain immune capacity (Gábor et al., 2020; Hystad et al., 2017). Oenocytes, though less directly involved in immunity, support lipid metabolism and detoxification (Lu et al., 2018). The activity of granulocytes and plasmatocytes is often reinforced by melanization, a biochemical response mediated by the prophenoloxidase cascade that leads to the deposition of melanin on invading microbes, including bacteria and fungi, thereby immobilizing and killing them (Hystad et al., 2017; Koleoglu et al., 2018; Zufelato et al., 2004).

Humoral immunity in honey bees involves the production of antimicrobial peptides (AMPs) such as defensin-1 and abaecin, which directly neutralize bacterial and fungal pathogens, while RNA interference provides antiviral defense (Danihlík et al., 2015; Yang et al., 2017). Together, these cellular and humoral mechanisms mentioned previously form a robust defense system. Still, they can be weakened by stressors such as *Varroa destructor* infestations, pesticide exposure, and infections caused by Deformed wing virus, the Gram-positive bacterium, *Paenibacillus larvae*, and the microsporidians, *Nosema* spp. (Al Naggar & Baer, 2019; Antúnez et al., 2009; Q. W. Chan et al., 2009; De La Mora et al., 2023; Koleoglu et al., 2018; Leponiemi et al., 2021; Sukkar et al., 2023).

Beyond their immune functions, hemocytes contribute to physiological homeostasis. They support nutrient transport and storage during periods of high metabolic demand, such as during larval development or adult foraging (Hystad et al., 2017; Negri et al., 2016). Hemocytes also regulate oxidative stress by producing reactive oxygen species (ROS) to combat pathogens while preventing excessive ROS accumulation that could harm host tissues (Koleoglu et al., 2018; Zufelato et al., 2004). In addition, they play a key role in wound healing by forming clots that prevent hemolymph loss and block pathogen entry (Hystad et al., 2017). These diverse functions highlight the central role of hemocytes in both immunity and overall honey bee resilience.

To investigate hemocyte contributions to honey bee immunity, we used clodronate liposomes (CLD) to deplete phagocytic hemocytes selectively. Clodronate liposomes, widely applied in arthropod models, serve as an effective tool for probing hemocyte function by inducing phagocytosis specifically in phagocytic cells. Once internalized, CLD is released and metabolized into a toxic ATP analog, which triggers programmed cell death (Frith et al., 1997). This approach has been successfully applied in *Drosophila melanogaster, Aedes aegypti, Amblyomma maculatum,* and *A. americanum* to assess the role of phagocytes in immunity and stress tolerance (Adegoke et al., 2024; 2023; Ramesh Kumar et al., 2021).

In this study, we applied CLD treatment to honey bees under optimal hive temperature conditions (32°C), reduced temperature conditions (22°C), and microbial infection. Our findings reveal that phagocytic hemocytes are essential for maintaining immune resilience and survival, particularly when bees face environmental stress and pathogen challenge. We demonstrate that CLD-mediated depletion of phagocytic hemocytes significantly compromises honey bee immune responses.

## MATERIALS AND METHODS

### Adult Honey Bee Collection

Worker bees used in the experiment were obtained from the USDA-ARS, Southeast Area Thad Cochran Horticultural Laboratory in Poplarville, MS, and the University of Southern Mississippi’s Lake Thoreau Environmental Center in Hattiesburg, MS. Frames containing capped brood close to emerging were pulled from the donor hives and placed in an incubator, where worker bees were allowed to emerge from the pupal stage at 32°C and 60% relative humidity (Alburaki et al., 2019). Newly emerged bees were collected from the brood frame and placed in cages provisioned with a 10-g pollen patty plug (Bee Pro Plus patties, Mann Lake, Minnesota, USA) and a 10 mL syringe containing 1:1 sterile sugar water (Alburaki et al., 2019). Following anesthetization and treatment injection, bees were sorted into their respective treatment groups and maintained in labeled cages supplied with fresh 10-g pollen patty plugs and sterile 1:1 sugar water, both of which were replaced every two days.

### Cage Experiment

The experimental groups were: (1) Untreated control, (2) Injection wound (Sham), (3) sterilized PBS injection (Vehicle control, pH 6.8), (4) CLD 1:5, and (5) control liposomes (LP) 1:5 (Standard Macrophage Depletion Kit, Encapsula Nano Sciences LLC, Brentwood, TN, USA). Clodronate-treated and control liposomes were diluted in the same PBS. A 1:5 dilution was selected because preliminary attempts with undiluted and 1:2 dilution resulted in high bee mortality by day 2 after exposure. Each treatment group consisted of three cages, with 30 bees per cage, for a total of 450 bees across all groups. For sampling, three bees were collected from each cage per day (technical replicates), yielding nine bees per group per day. Hemocyte counts were acquired from a total of 45 bees per day across all treatment groups.

Immediately post-injection, bees were transferred to new cages corresponding to their treatment group and placed in an incubator at 32°C for 30 minutes to recover. Each cage was provisioned with a 10-g pollen patty plug and a syringe containing 1:1 sterile sugar water, replaced every two days. Survival was monitored daily, with dead bees counted and removed. Bees used in the 32°C and 22°C experiments were 5 days old at the time of injection. Fifteen-day-old bees were used in later experiments to approximate forager age, while 5-day-old bees represented nurse age. Younger bees (1-3 days old) were excluded because injection trials at this age resulted in high mortality, whereas 4-5-day-old bees tolerated the procedure. Incubator conditions were maintained at 32°C and 60% RH, while 22°C trials were conducted under ambient conditions. PBS and CLD groups were the primary comparison, as CLD was the experimental focus.

### Chemical Depletion of Phagocytic Hemocytes

The experiment utilized CLD injection as a targeted tool to chemically deplete phagocytic hemocytes. The specific purpose was to induce immunosuppression to study the dynamics of hemocyte loss and recovery, as measured by changes in total and differential hemocyte counts, and to determine the effect of depletion on bee survival. For injections, five bees were simultaneously anesthetized by submersion in ice and then secured at the head and thorax with scotch tape on a chilled Petri dish to maintain anesthesia. Injections were administered with a Hamilton syringe fitted with a 33-gauge needle (Hamilton Company, Franklin, MA, USA). The treatment solution (0.1 µL) was injected between the 2^nd^ and 3^rd^ abdominal tergites, offset laterally to avoid vital organs (Koleoglu et al., 2017, 2018). After injection, bees were placed into cages according to their treatment group. The needle was sterilized in 70% ethanol between injections and allowed to dry before use.

### Hemolymph Extraction

To determine the optimal extraction site, hemolymph was sampled from the base of the head at removed antennae sections, abdominal punctures, and decapitation wounds. Honey bees were first anesthetized on ice. For antennal hemolymph collection, bees were surface sterilized with 70% ethanol, and residual ethanol was removed with a Kimwipe before evaporation. Antennae were removed with forceps, eliciting a small droplet of hemolymph that was collected with a pipette. Gentle pressure was applied to the thorax to increase hemolymph yield (Borsuk et al., 2017). For hemolymph that was collected at the site of decapitation, the head was removed, and thoracic pressure was applied to pool hemolymph at the neck region for pipette collection (Borsuk et al., 2017). For abdominal collection, bees were punctured in the abdomen, and rhythmic thoracic pressure stimulated hemolymph flow. Samples with visible debris or discoloration suggesting contamination were discarded (Borsuk et al., 2017). After comparing methods, the antennal approach was used for subsequent experiments due to its higher hemolymph yield and reduced contamination risk. All hemolymph was suspended in sterile phosphate-buffered saline (PBS, pH 6.8) and kept on ice to preserve cellular integrity. Hemolymph volumes were quantified for each bee individually.

### Hemocyte Quantification

Hemolymph was collected from three bees from each cage for a total of nine bees per treatment group for 5-day-old and 15-day-old bees. Hemocyte counts were performed using hemolymph collected from the antennae of individual honey bees. For each bee, 2.5 µL of hemolymph was diluted in 2.5 µL of sterile PBS (pH 6.8) using a 10 µL pipette (Dostálková et al., 2020). Hemolymph samples were stained with 0.4% trypan blue and counts were obtained using an iNcyto C-Chip Neubauer hemocytometer with calculations performed according to the manufacturer’s standard formula (Adegoke et al., 2023). Differential counts of granulocytes, plasmatocytes, prohemocytes, and oenocytes were also recorded. Hemocyte counts were measured on days 1, 2, 4, and 7 post-injections. Bee age at injection was recorded as baseline, and subsequent days reflected age progression during the experiment. In temperature trials (32°C vs 22°C) and bacterial challenge assays (*E. coli*, *S. aureus*, and control), 45 hemocyte counts were performed per day, totaling 180 counts per 7-day run. Across five 7-day runs, 900 hemocyte counts were conducted. For survival analysis, cages were monitored daily over 7 days, and survival percentages were used to assess the treatment effect.

### Bacterial Challenge

*Escherichia coli* strain DH5-alpha and *Staphylococcus aureus* strain RN4220 were extracted from individual plate colonies. Each colony was inoculated into 10 mL of medium: Luria-Bertani broth (LB) for *E. coli* and Tryptic Soy broth (TSB) for *S. aureus* suspension (Adegoke et al., 2023). Cultures were incubated overnight at 37°C on a rocker. The following day, live bacteria were collected from the culture supernatant, quantified by spectrophotometry at OD600, and diluted to 4 × 10^6^ CFU/mL in 1:2 sugar-to-water (Kešnerová et al., 2017). After receiving the assigned treatments, honey bees were deprived of their sterile 1:1 sugar water for 6 hours and then given the bacterial suspension for 24 hours.

### Visualization of Phagocytosis Using Fluorescent Beads: *In vivo* Assay

Phagocytic activity was assessed using green fluorescently conjugated carboxylate beads (2 µM; Thermo Fisher Scientific, Waltham, MA, USA) (Adegoke et al., 2023). Honey bees were anesthetized on ice and injected with 1 µL of a solution containing diluted fluorescent beads (1:5 in buffer) and 0.75 mM CM-DiL in sodium citrate anticoagulant buffer (pH 7.4). Injections were performed between the 2^nd^ and 3^rd^ tergites (Hystad et al., 2017). Following injection, bees were maintained in an incubator at 32°C and 60% RH for 30 minutes to recover. Hemolymph was then collected from anesthetized bees and stained for visualization by confocal microscopy. Both beads and liposomes used in this assay measured 2 µm in diameter. Representative images are provided in the text and supplementary materials.

### Hemocyte Staining

Hemolymph samples were diluted 1:1 in sterile PBS (pH 6.8), and 20 µL of the mixture was pipetted onto coverslips. Samples were incubated at 32°C for 40 minutes to allow hemocyte adhesion, then fixed with 4% paraformaldehyde for 20 minutes. Cells were permeabilized with 0.5% Triton X-100, followed by three PBS washes (Adegoke et al., 2023). For nuclear staining, 15 µL of Vectashield mounting medium containing DAPI was applied, and coverslips were sealed with nail polish for long-term storage at 4°C. For membrane staining, Vybrant CM-DiL (0.75 mM in PBS) was applied following the injection protocol described above (Hystad et al., 2017). CM-DiL is a lipophilic dye commonly used for cell tracking and invertebrate hemocyte staining. Phagocytosis of fluorescent beads by hemocytes was assessed by confocal microscopy to evaluate immune activity.

### Image Acquisition

Fluorescently stained hemocytes were imaged using a Leica STELLARIS STED (Leica Microsystems, Wetzlar, Germany) with either a 63X or 100X objective. Z-stacks were acquired to capture the full depth of hemocytes. Excitation was performed using wavelength-specific lasers: 405 nm for DAPI, 488 nm for phalloidin, 505 nm for fluorescent beads, and 545 nm for CM-Dil. Brightfield images were captured simultaneously using the secondary fluorescent laser channel, with DAPI as the primary channel. Post-acquisition adjustments to light exposure were performed using Leica’s built-in processing software. Final image panels were assembled in PowerPoint for analysis and visualization.

### Data Analysis

To control cage effects and assess consistency among biological replicates, three replicate cages were maintained for each treatment and control group. Bees subsampled for hemolymph collection were excluded from survival analyses. Figures and statistical analyses were performed using R Studio (Version 2024.12.1+563) and GraphPad Prism (Version 10.4.1), except hemocyte images. Figures were generated utilizing multiple R libraries, mainly “ggplot2”, “doby”, “plyr”, “performanceAnalytics”, and “data.table”. For line graphs, error bars represent the standard deviation (SD) unless stated differently in the figure’s caption. At the same time, boxplots display the median, first and third quartiles, and both maximum and minimum values of variables. All datasets were tested for normality using the Shapiro test before conducting statistical analyses. The non-parametric Kruskal–Wallis rank test was conducted at a 95% confidence interval with three levels of significance (*p* < 0.05, < 0.001, < 0.001) on data that failed the normality test. Multiple comparisons and p-values were adjusted with the Benjamini–Hochberg method when applicable, as the data failed the normality test. Few datasets were normally distributed; in these cases, one-way ANOVA with Tukey *ad hoc* test was used to assess group statistical differences. Survival analyses were conducted using both the Kaplan-Meier model and simple overall averages of dead bees per treatment. Additionally, heatmaps were generated using the “pheatmap” package to assess the daily and overall hemocyte variance. Variance was analyzed across distinct treatments (injection type: control, wound, PBS, CLD, LP) and conditions (temperature for 5-day-old cohorts; bacterial challenge for 15-day-old cohorts). Generalized Linear Mixed-effect Models (GLMM) were used to evaluate and predict the impact of the treatments and bacterial challenge on hemocyte count. The core model is as follows: (GLMM <-response variable (Hemocytes per mL) ∼ predictor variables (Treatment + Hemocyte Cell Types; or Treatment alone), data = x, family = quasi). The control category was deliberately chosen to be reflected in the model’s intercept for easier handling of the model’s estimation values. GLMM analyses were separately conducted on three different experimental groups: A) 5-day-old bees at 32°C, B) 5-day-old bees at 22°C, and C) 15-day-old bees at 22°C.

## RESULTS

### Treatment Effect on Honey Bee Survival

A significant correlation was observed between hemocyte depletion via CLD/LP treatment and reduced survival in honey bees at both 32°C and 22°C (*p* < 0.001), suggesting the importance of hemocytes in sustaining survival across temperatures (Fig. 2A, 2B). The experiment at 32°C was ended on day 4 for all groups due to the high mortality seen in all injection groups with the most deaths occurring in the CLD group leaving 5 bees left to sample. Similar patterns of reduced survival were observed in PBS and LP-injected groups, whereas the control group showed the highest survival rates (*p* < 0.001; Fig. 2A, 2B). At 22°C, a sufficient number of bees survived through the 7-day trial for all treatment groups for hemocyte analysis, although survival was still significantly decreased (*p* < 0.001) in PBS and CLD-treated groups compared to controls (Fig. 2A, 2B). While the Kaplan-Meier model provides better prediction and accuracy, the overall average bee mortality showed the same trends with significantly lower mortality in the control group compared to other treatment groups at 32°C (*p* < 0.05) and 22°C (*p* < 0.01) (Fig. 2A, 2B).

**Figure 1:**
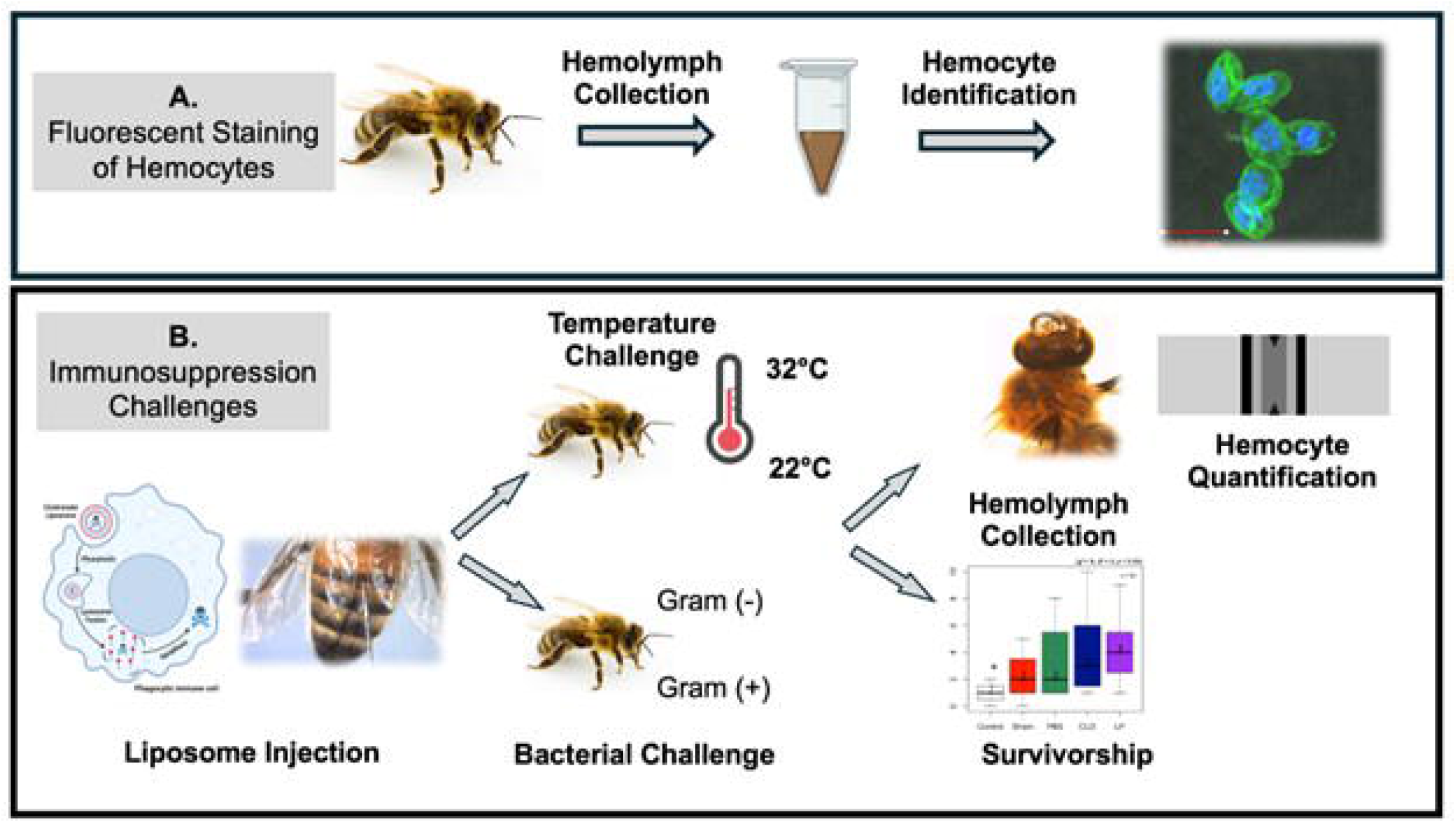
Graphical abstract: **(A)** Identification of hemocyte types in adult honey bee hemolymph. Honey bee hemolymph was extracted and fluorescently stained to distinguish between the four hemocyte types in circulation. **(B)** Effects of temperature or bacterial challenge on hemocyte populations and survival of adult bees with reduced circulating hemocytes. Bees were anesthetized and injected with liposomal immunosuppressants (clodronate liposomes) to test the effects of reduced circulating hemocyte populations on honey bees. Further challenges, such as varying ambient temperature or bacterial challenges, were conducted after reducing circulating hemocyte populations in the later experimental runs. Bee mortality was monitored daily to assess the treatment effects. Hemolymph and total/differential hemocytes were analyzed to observe changes in circulating hemocyte populations and bee mortality post-injection.

**Figure 2.**
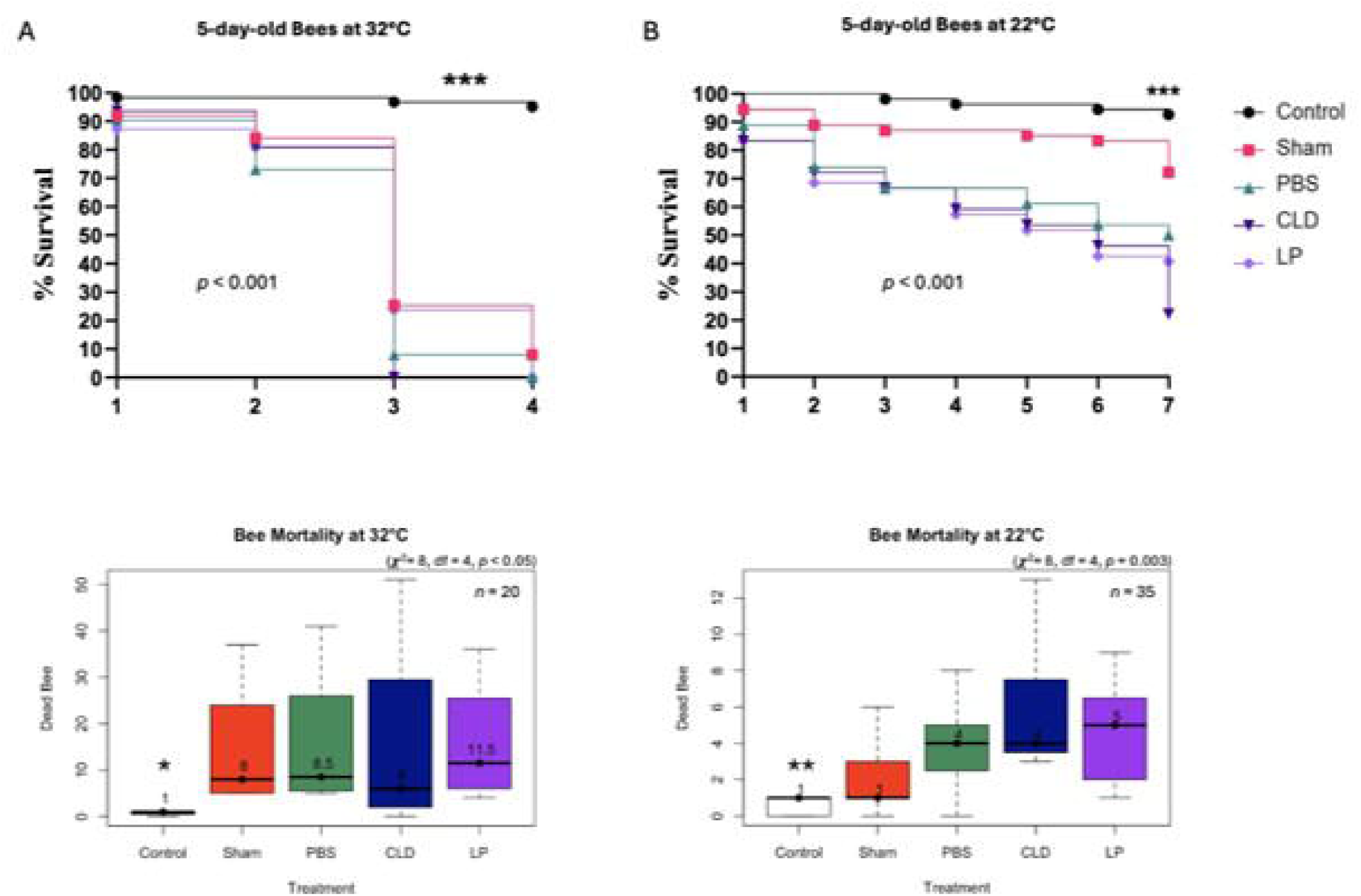
Effect of treatment on the survival of 5-12-day-old bees at 32°C and 22°C. Kaplan-Meier survival curves of bees kept in cages at 32°C (A) and 22°C (B) and monitored daily for 7 days. Significance was determined through the model log-rank test for survival curves, and the Kruskal-Wallis non-parametric test was used to assess statistical differences between boxplot groups (A and B). Median is displayed for each boxplot, and asterisks mark the levels of significance (*p* < 0.05*, *p* < 0.01**, *p* < 0.001***).

### Treatment Effect on Hemolymph Volume

Hemolymph volume was also significantly affected by temperature and CLD treatment. At 32°C, CLD-treated bees showed significant reductions in hemolymph volume on all sampling days (*p* < 0.001) compared to the Control group, while at 22°C, hemolymph volume was only significantly lower on day 1 (*p* < 0.01; Fig. S1A, B). CLD-treated bees at 22°C retained higher overall hemolymph volumes than those at 32°C, with averages of 2.7 µL and 3.5 µL, respectively (Fig. S2). In contrast, PBS-treated bees at 32°C maintained slightly higher volumes than their 22°C counterparts (Fig. S2A, B). Average control hemolymph volumes at 22°C and 32°C significantly (*p <* 0.001) exceeded those of treatment groups (Fig. S1A, B). Overall, circulating hemocyte counts correlated with hemolymph volume (Fig. S1, S2).

### Treatment Effects on Hemocyte Count

Total hemocyte counts further highlighted temperature-dependent effects 5-day old bees. At 32°C, PBS and CLD-treated bees showed significant reductions on days 1, 2, and 4 (*p* < 0.001, *p* < 0.001, and *p* < 0.01, respectively; Fig. 3A). At 22°C, significant reductions occurred on days 2, 4, and 7 (*p* < 0.05, Fig. 3A). Initial hemocyte counts in control bees were higher at 32°C (∼1,426,000 hemocytes/mL) compared to 22°C bees (∼1,000,000 H/mL). By the end of the experiment, controls at 32°C retained ∼1,298,000 H/mL, whereas controls at 22°C declined to ∼740,000 H/mL, suggesting reduced hemocyte population retention at lower temperatures (Fig. 3A). Total hemocyte abundance between temperatures reveals no significant difference between 22°C and 32°C in 5-day-old bees (Fig. 4A). LP-injected bees maintained total hemocyte counts similar to those of CLD-treated bees across both conditions (Fig. 4).

**Figure 3.**
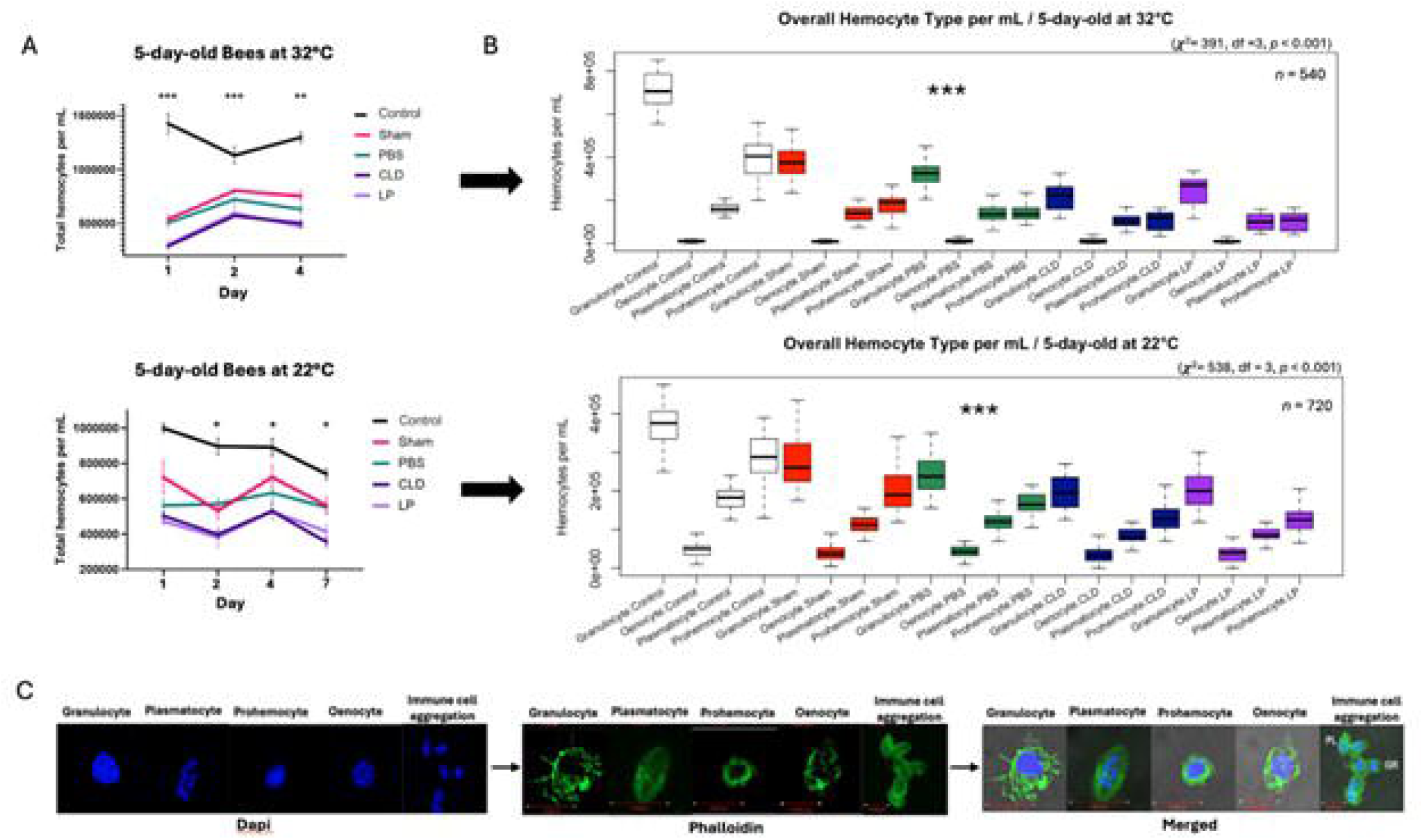
Effect of treatment on hemocyte abundance and characterization in 5-day-old bees. Total hemocyte count over time (A) and differential hemocyte counts by cell type (B). Fluorescent confocal microscopy images of major hemocyte subpopulations (C). Statistical differences between groups were determined using the Kruskal-Wallis test. Median is displayed for each boxplot, and asterisks mark the levels of significance (*p* < 0.05*, *p* < 0.01**, *p* < 0.001***). Error bars in line graphs represent the standard deviation (SD). Abbreviations: PL, plasmatocytes; GR, granulocytes. Scale bar = 10µm.

**Figure 4.**
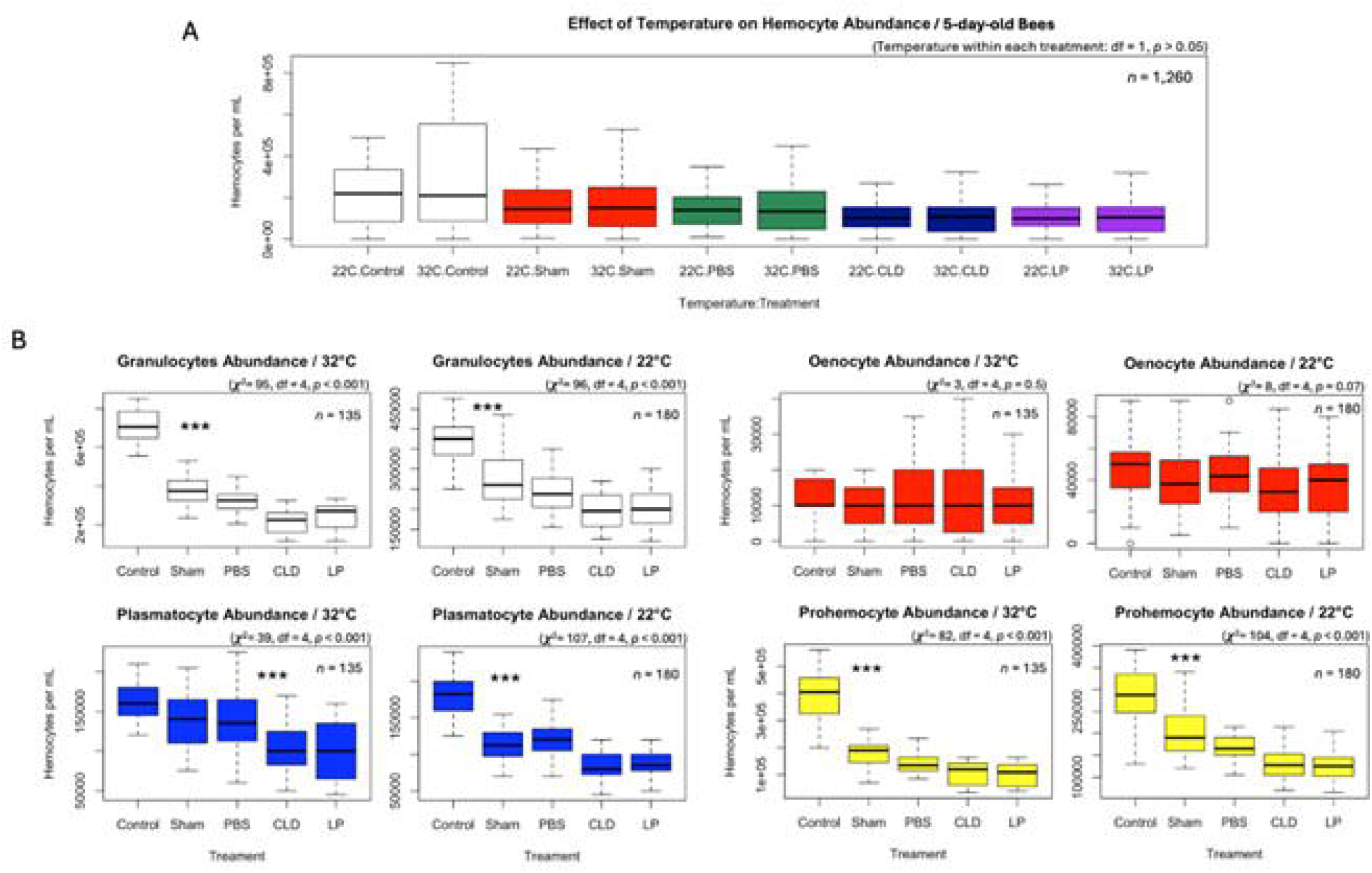
Effect of temperature on hemocyte abundance in injected 5-day-old bees. Overall (A) and differential (B) hemocyte abundance across treatments and temperatures. Kruskal-Wallis test was used to assess statistical differences among groups, and asterisks mark the levels of significance (*p* < 0.05*, *p* < 0.01**, *p* < 0.001***).

Differential hemocyte counts revealed specific cell type responses to treatment and temperature. At 22°C and 32°C, CLD-treated bees exhibited significantly reduced (*p* < 0.001) granulocyte, plasmatocyte, and prohemocyte abundance (*p* < 0.05; Fig. 3B, 4B). Oenocyte counts remained stable across treatments and temperatures, with no significant trends (Fig. 3B, 4B). When comparing control groups between temperatures, granulocytes showed the largest differences in population, followed by prohemocytes and plasmatocytes, respectively. Similar trends were observed under CLD treatment, highlighting granulocytes as the hemocyte type most reduced in circulating populations under temperature change (Fig. 3B, 4B).

### Hemocyte Variation by Age

The injection treatments were repeated with 15-day-old bees. Survival analysis revealed that 15-day-old bees exhibited significantly greater tolerance to hemocyte depletion compared with 5-day-old bees (*p* < 0.001; Fig. 2, 5A). In the CLD-treated 15-day-old cohort, survival declined to 46%, whereas the 5-day-old CLD-treated group showed substantially lower 22% survival under comparable conditions (Fig. 2B, 5A). On average, the control group had significantly less death compared to the other treatments (*p <* 0.05) (Fig. 5A). With the addition of a bacterial challenge of either *E. coli* or *S. aureus* paired with the injection treatments, the survival decreased further in comparison to injection treatments not challenged with bacteria but this difference was not significant (Fig. 5A-C).

**Figure 5.**
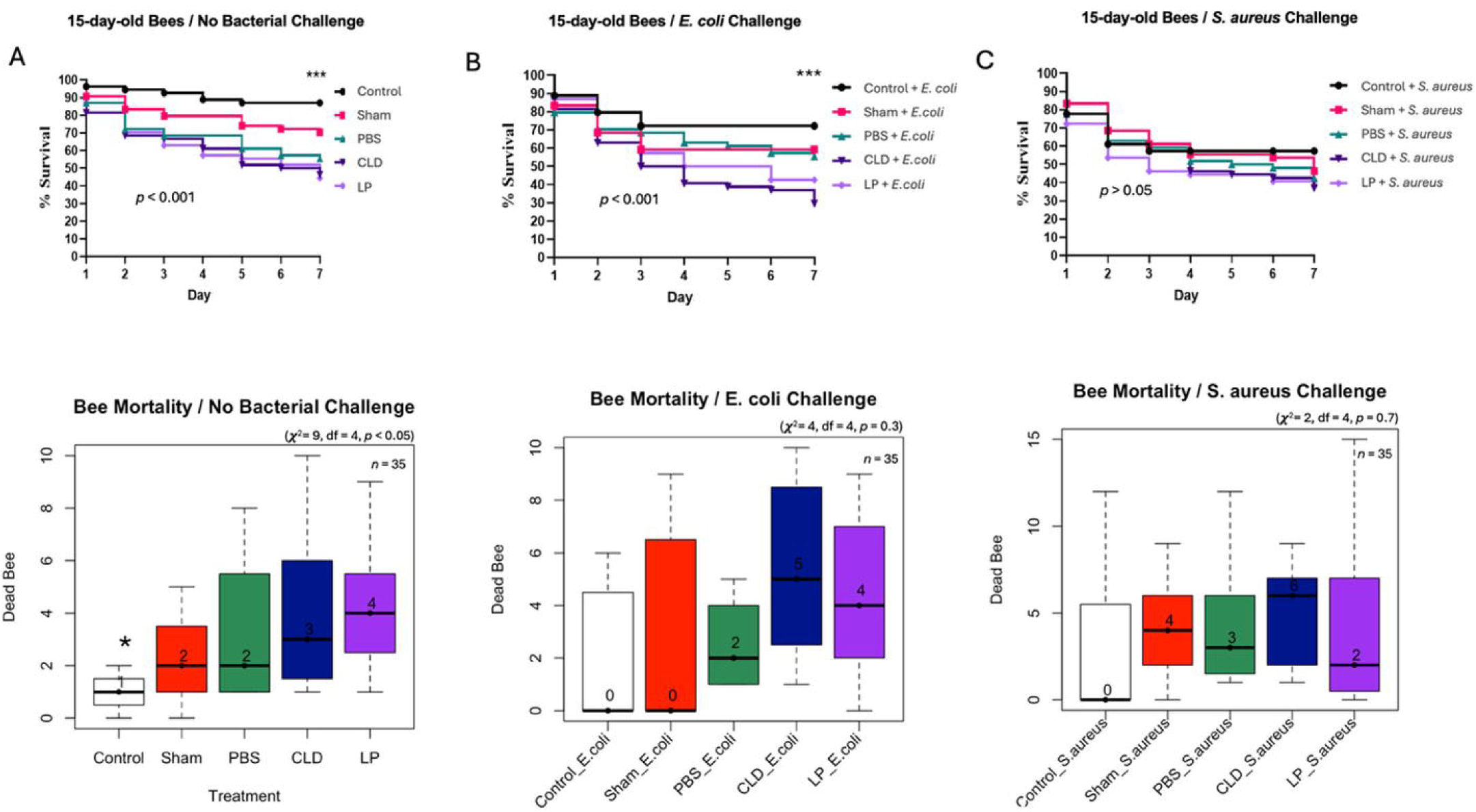
The effect of treatment and bacterial challenge on the survival of 15-day-old bees. Daily and overall survival of treatment-injected bees without a bacterial challenge (A), and with a subsequent challenge of *E. coli* (B), or *S. aureus* challenge (C). The Levels of significance are (*p* < 0.05*, *p* < 0.01**, *p* < 0.001***), determined through the Kaplan-Meier survival model for the line graphs and the Kruskal-Wallis test for the box and whisker plots.

Comparison of treatment with bacterial challenge helps to reveal the necessity of hemocytes in combating infection and the rate at which bacterial challenges reduce hemocyte stress in the immune system. The total hemocyte abundance of the unchallenged 15-day-old bees revealed significantly more hemocytes compared to the bacterial challenge groups (*p <* 0.01, *p <* 0.001) (Fig. 6A, 6C). Differential hemocyte abundance revealed a similar trend to 5-day old bees. In the CLD and LP groups granulocytes were negatively affected the most with effects further intensified by bacterial challenges (Fig. 6B). Similarly, prohemocytes and plasmatocytes showed the second and third greatest reductions, respectively (Fig. 6B).

**Figure 6.**
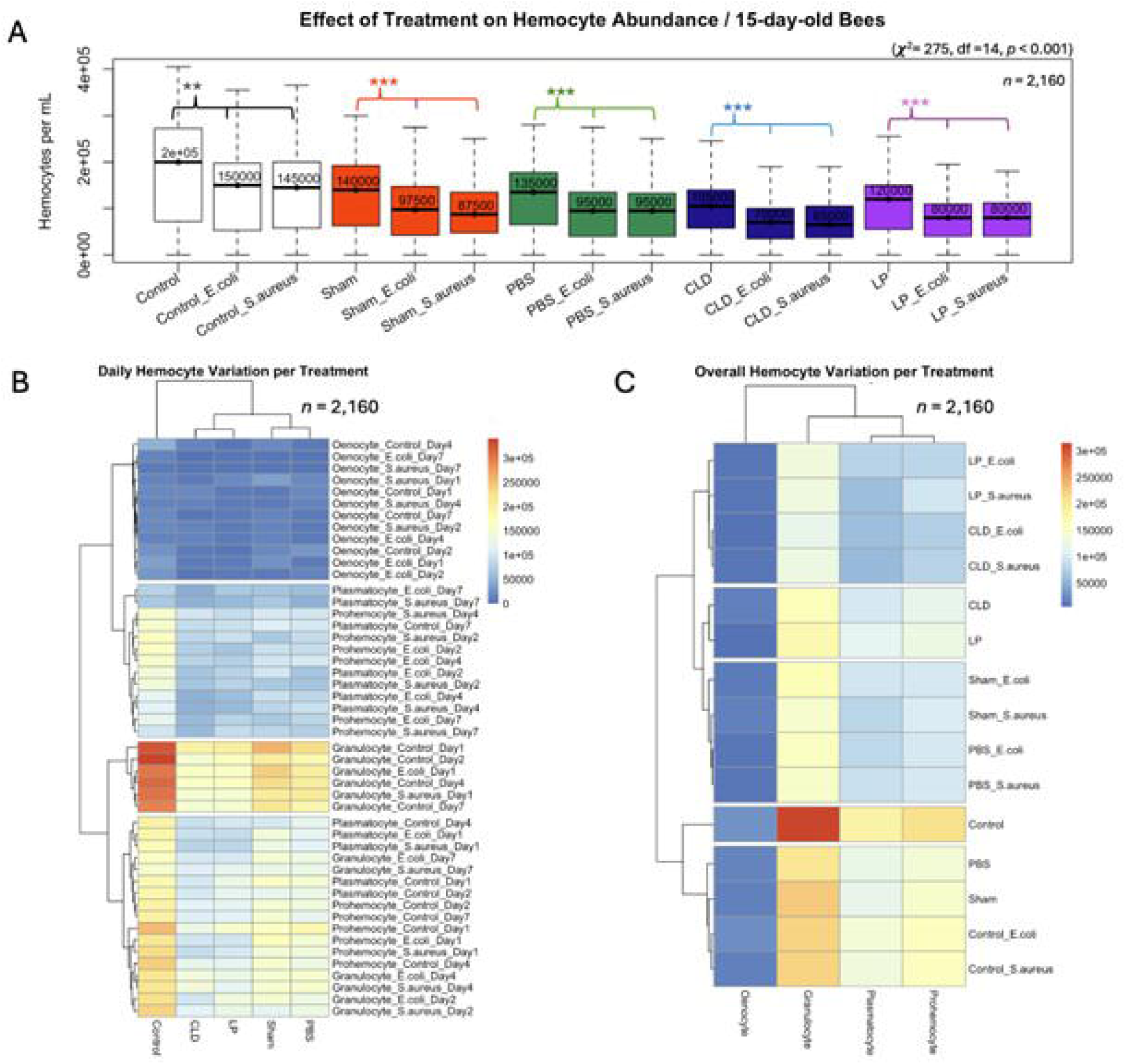
Overall effect of treatment on hemocyte abundance in 15-day-old bees. Overall average of hemocyte per treatment (A), heatmap visualization of hemocyte variations vis-à-vis treatment, cell type, and day (B), overall hemocyte count per treatment only (C). Kruskal-Wallis non-parametric test was conducted to assess the hemocyte variation vis-a-vis the treatment, with asterisks denoting the levels of significance (*p* < 0.05*, *p* < 0.01**, *p* < 0.001***) between groups.

The total hemocyte abundance was mapped to display the trend of decreased cellular immunity over the course of age in honey bees. Bee nurses had significantly higher hemocyte abundance in comparison to foragers and treatment injection groups (*p <* 0.001) (Fig. 8A). The foragers control group retained significantly higher overall hemocyte abundance (*p <* 0.001) in comparison to the treatment injection and bacterial challenge groups, with few signs of recovery. The overall hemocyte counts showed that nurse bees retained significantly (*p <* 0.001) higher hemocyte abundance (161,434 H/mL) compared to forager bees (105,916 H/mL), Fig. 7.

**Figure 7.**
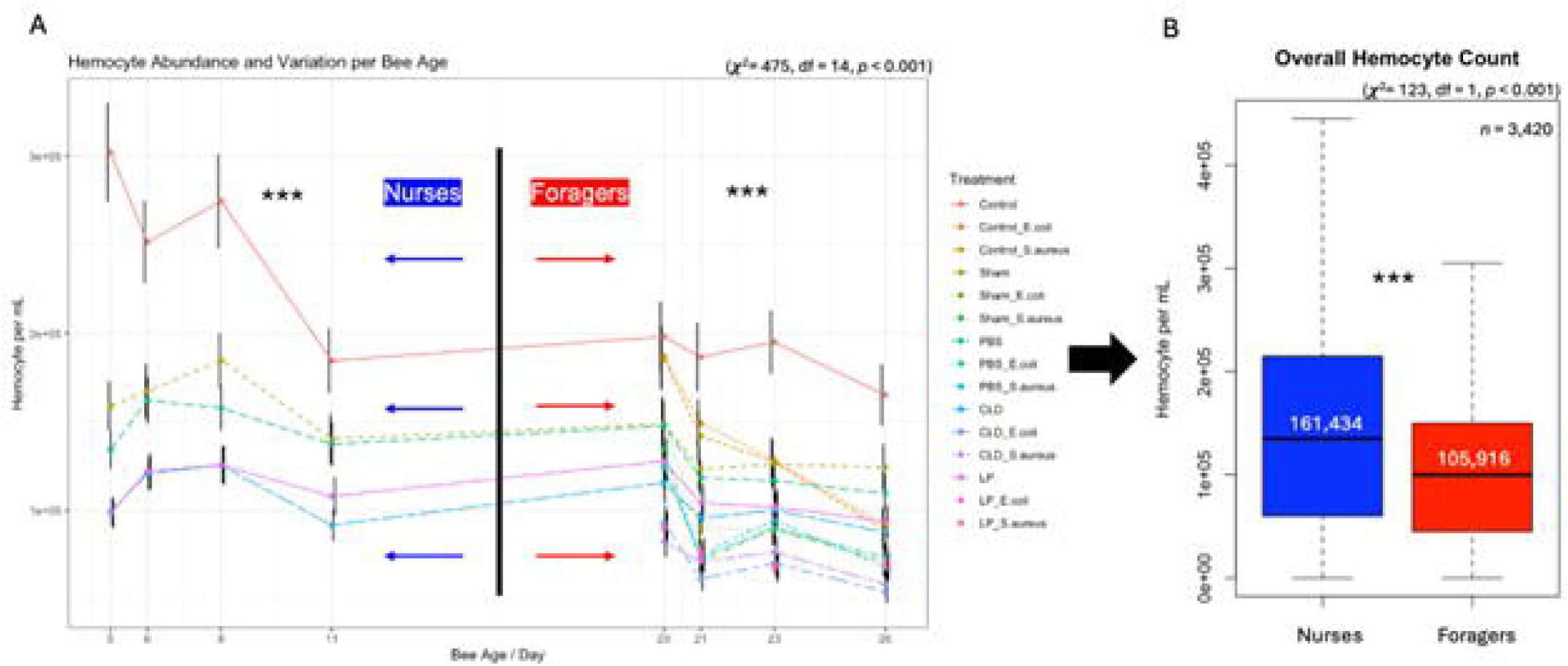
Longitudinal and overall analyses of hemocyte variations by bee age, treatment, and bacterial challenge. (A) line graph of hemocyte count for both bee age groups (5-day-old “Nurses”, and 15-day-old “Foragers”), (B) overall averages sorted by bee ages (Nurses vs. Foragers). Error bars in the line graph represent the standard error (SE). Median is displayed in each boxplot. Kruskal-Wallis non-parametric test was conducted to assess the hemocyte variation vis-a-vis the treatment, with asterisks denoting the levels of significance (*p* < 0.05*, *p* < 0.01**, *p* < 0.001***) between groups.

## DISCUSSION

### Bee Hemocyte and Survival Responses to Clodronate

Consistent with previous reports, we observed morphological and functional heterogeneity in hemocytes from bees sampled between 5-12 days of age and 15-22 days of age (Gábor et al., 2020; Hystad et al., 2017). CLD injections significantly reduced total hemocyte populations, with granulocytes and prohemocytes as the primary targets. Correspondingly, hemolymph volume was also reduced. Comparisons with previous CLD studies in *D.*, *A. aegypti, A. maculatum*, and *A. americanum* revealed similar findings (Adegoke et al., 2023, 2024; Kumar et al., 2021). Like these arthropods, honey bees exhibited decreased granulocyte counts and reduced survival following CLD treatment. However, honey bees responded more negatively to control liposome injections than other arthropods studied, which was correlated with immunosuppression and reduced survival. This response indicates a heightened sensitivity to liposomal treatments and underscores the vulnerability of honey bee cellular immune defenses.

The distinct survival and hemocyte responses of honey bees compared with solitary or free-living arthropods may be linked to their social lifestyle. Honey bees employ both individual and collective immune defenses (social immunity), potentially reducing selective pressure on individual immune functions (Baxter et al., 2017; Cremer et al., 2007; Evans & Spivak, 2010; Meunier, 2015; Rosche et al., 2021). As a result, interventions targeting hemocytes may disproportionately impair survival in honey bees compared with solitary species. Further research should explore the interaction between social immunity and hemocyte-mediated defenses to clarify how these mechanisms jointly shape honey bee resilience (Simone et al., 2009). Expanding CLD studies to insects across a spectrum of sociality, from solitary, primitively social, and eusocial, would provide a comparative framework to identify conserved versus divergent immune patterns associated with sociality. Such an approach could help disentangle the relative contributions of individual and collective immune defenses, with broader implications for pollinator and insect health.

### Effects of Temperature and Hemocyte Loss on Young Bees

Honey bee colonies typically maintain brood-rearing temperatures at 32-36°C (Kutby et al., 2024; Southwick & Heldmaier, 1987; Tautz et al., 2003). We observed that bees injected with CLD at 5 days old experienced near total mortality by day 4 when maintained at 32°C after injection (Fig. 2A). In contrast, bees maintained at 22°C survived at higher rates, with ∼20% greater survival in CLD-treated groups (Fig. 2A, B). These findings suggest that while 32°C supports development of healthy bees, it becomes detrimental under conditions of hemocyte depletion, likely due to elevated metabolic demands (Hystad et al., 2017; Marringa et al., 2014; Morimoto et al., 2011).

At 32°C, accelerated metabolism may increase hemocyte turnover and exacerbate CLD-induced apoptosis, contributing to sharp declines in hemolymph volume (2.7 µL; Fig. S1A). Conversely, CLD treated bees maintained at 22°C retained greater hemolymph volume (3.5 µL; Fig. S1B), consistent with enhanced hemolymph retention under cooler conditions. Differential hemocyte counts also indicated that prohemocytes and oenocytes declined more gradually at 22°C (Fig. 3A-B), suggesting a potential for more efficient replenishment when metabolism is restrained (Glass & Harrison, 2024; Kovac et al., 2007; Stupski & Schilder, 2021). These results align with studies showing that suboptimal temperatures can enhance insect stress tolerance by downregulating energy-intensive processes (MacMillan et al., 2017; Shan et al., 2024). Although 22°C can be physiologically stressful for nurse bees, reduced metabolic activity at this temperature appears to mitigate the costs of hemocyte loss, buffering survival during immune collapse (Ellis & Rangel, 2024; Ulgezen et al., 2024).

### Age-Specific Response to Hemocyte Population Reduction

Fifteen-day-old honey bees exhibited greater resilience to CLD hemocyte depletion than 5-day-old bees, with survival rates of ∼46% compared to ∼22% in younger cohorts (Fig. 2B, 5A). This survival advantage occurred despite lower baseline hemocyte counts in older bees (161,434 H/mL for nurses vs. 105,916 H/mL for foragers in; Fig. 2B, 5A, 7), suggesting an age-dependent tradeoff between cellular immunity and alternative survival mechanisms (Gábor et al., 2020; Hystad et al., 2017; Ravaiano et al., 2018; Schmid et al., 2008; Yelkovan et al., 2021).

Younger bees, which invest heavily in cellular defenses during early adult stages, maintained higher granulocyte levels (346,666 H/mL vs. 298,300 H/mL in 15-day-olds; Fig. 3B, 4B) but were more vulnerable when hemocyte populations were reduced. In contrast, older bees, despite diminished hemocyte counts, may rely on compensatory humoral or physiological stress adaptations such as AMP, prophenoloxidase, and antioxidant production peaking day 28 in age according to literature (Jefferson et al., 2013; Kunat-Budzyńska et al., 2025; Schmid et al., 2008). Granulocytes, prohemocytes, and plasmatocytes exhibited similar trends of reduction in 15-day-old bees compared to 5-day-old bees. These results are consistent with the hypothesis that aging workers shift from hemocyte-mediated immunity to broader stress tolerance as they transition to foraging roles (Bull et al., 2012; Hystad et al., 2017; Schmid et al., 2008).

### Infection Susceptibility in Hemocyte-Deficient Bees

Bacterial infections in 15-day-old bees exacerbated hemocyte depletion and survival rate, consistent with clodronate liposome studies in other arthropods (Adegoke et al., 2024; Ramesh Kumar et al., 2021). Among hemocyte types, granulocytes were most susceptible to challenge with *E. coli* or *S. aureus*, followed by prohemocytes, whereas plasmatocytes were comparatively resilient (Fig. 6B, C). This hierarchy likely reflects functional specialization; granulocytes act as primary phagocytes and are highly vulnerable to immunosuppression (Adegoke et al., 2023, 2024; Hystad et al., 2017; Ramesh Kumar et al., 2021), plasmatocytes contribute primarily to encapsulation and wound healing, and may be less reliant on rapid pathogen recognition (Gábor et al., 2020; Hystad et al., 2017), while prohemocyte depletion may result in differentiation of prohemocytes into granulocytes in the advent of immunosuppression (Hystad et al., 2017; Marringa et al., 2014). Even at 22°C, where reduced metabolism buffered CLD-induced mortality, bacterial infections drove hemocyte populations toward critical thresholds ∼220,000 H/mL by day 7 (Table. S1).

### Intrinsic Bee Immune Constraints

Across experiments, a suggested critical hemocyte threshold of ∼220,000 H/mL was identified, below which survival was not sustained. Bacterial challenges accelerated depletion to this level, with *E. coli* reducing counts to 216,666 H/mL (Fig. 6, 7). This threshold likely defines the minimum circulating hemocyte population necessary for immune competence and survival. Honey bees’ reliance on mitotic division of circulating hemocytes, rather than dedicated hematopoietic organs as in *Drosophila*, limits their capacity to recover from hemocyte depletion (Marringa et al., 2014; Richardson et al., 2018). This constraint may render bees particularly vulnerable to combined stressors, including pathogens, pesticides, and environmental challenges, with direct consequences for colony health. The limited regenerative potential of their immune system underscores the importance of maintaining colony-level protections, such as social immunity and environmental stability, to support pollinator resilience and sustain critical ecosystem services.

## Conclusion

This study demonstrates that phagocytic hemocyte abundance is strongly correlated with honey bee survival. Selective depletion of these cells using CLD revealed that bees maintained at optimal hive temperatures (32°C) experienced near-total mortality, whereas those at suboptimal temperatures (22°C) showed reduced mortality likely due to reduced metabolic demands. Age also influenced resilience: 15-day-old bees survived better than 5-day-old bees despite lower baseline hemocyte counts, suggesting compensatory physiological or behavioral mechanisms. Bacterial infections (*E. coli* and *S. aureus*) further exacerbated hemocyte depletion and mortality, highlighting the compounded impact of multiple stressors. A critical hemocyte threshold of ∼220,000 H/mL was identified, below which survival became unsustainable, underscoring the limited regenerative capacity of honey bee hemocytes under chronic stress.

These findings emphasize the importance of maintaining robust hemocyte populations for individual and colony health. Future work should explore strategies to enhance hemocyte resilience through nutritional support, selective breeding, or other interventions to strengthen honey bee immunity and mitigate the effects of environmental and pathogenic pressures.

### Technical Limitations

Several technical limitations should be considered in this study. First, hemocyte counts may vary with the collection method, and our consistent use of the other antennae method may not account for localized hemocyte distribution. Second, CLD’s effects may extend beyond the intended phagocytic cells, potentially impacting other processes. Third, experiments were limited to two temperatures (22 and 32°C) and two age groups (5- and 15-day-old bees). Expanding temperature and age ranges would offer a more complete picture of immune resilience. Using a fixed bacterial concentration does not replicate natural infection dynamics. Varying pathogen loads and types would be more informative. Future work should include transcriptomic or proteomic analyses. Finally, laboratory conditions lack social immunity factors, where field studies are necessary to validate findings in the natural hive context.

**Table 1.**
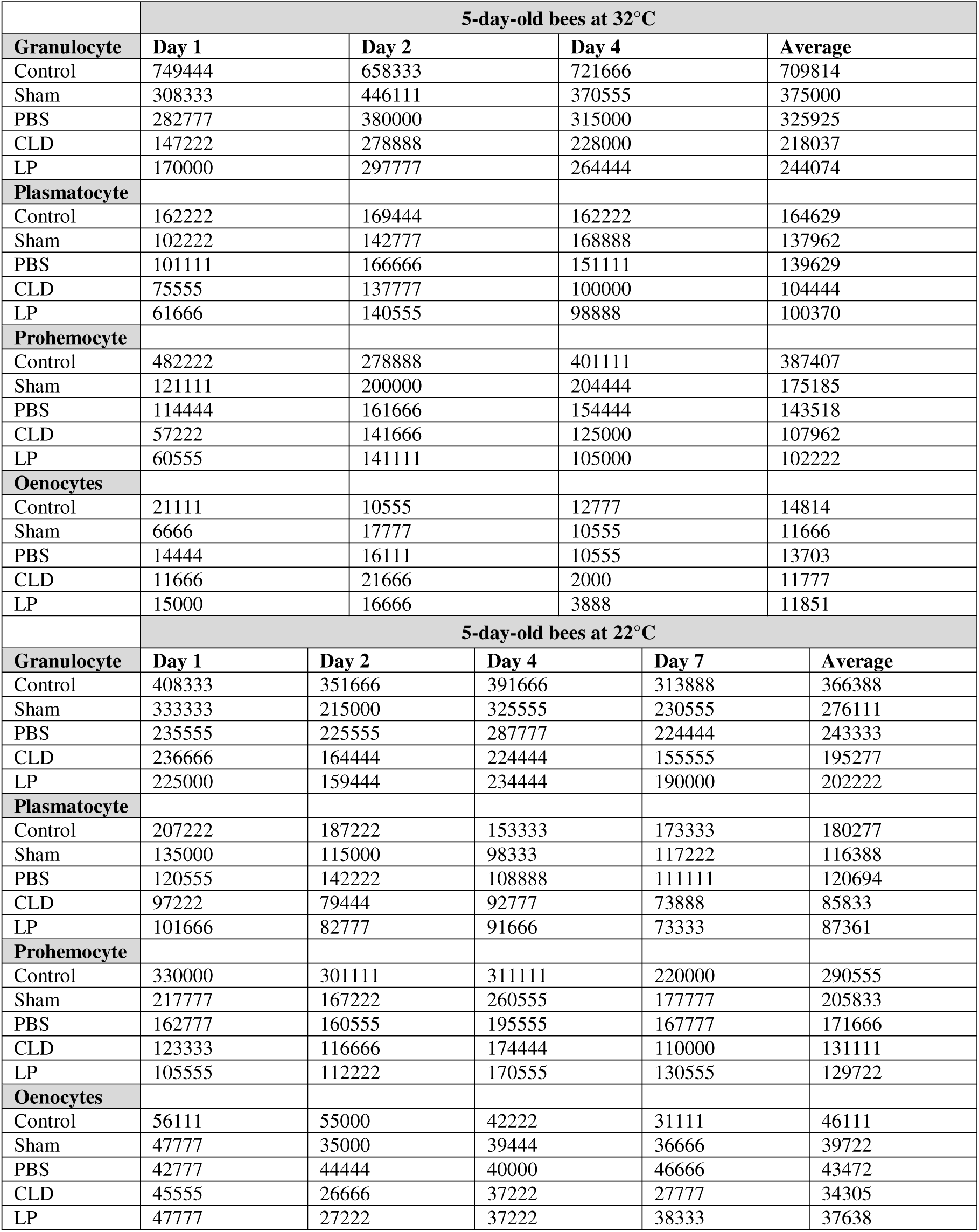
Summary of hemocyte cell count per mL displayed by daily average and total average. Hemocyte cell counts were conducted on 5-day-old bees kept at both 32 °C and 22°C.

**Table 2.** Generalized Linear Model (GLMM) with mixed effects, predicting the effects of treatment on hemocyte quantity and cell type. The model was conducted on hemocytes per mL data as a response variable and both treatment and hemocyte cell type as predictor variables: (GLMM <-response variable ∼ Treatment + Type, data = x, family = quasi), for 5-day-old bees at 32 and 22°C separately. Levels of significance are *p* < □ 0.001***.

| GLMM model output / Hemocyte of 5-day-old bees at 32°C |  |  |  |  |
| --- | --- | --- | --- | --- |
|  | Estimate | Std Error | T-Value | P-Value |
| (Intercept) | 521342 | 10672 | 48.85 | < 0.001*** |
| CLD | -209167 | 12368 | -16.91 | < 0.001*** |
| LP | -204537 | 11864 | -17.24 | < 0.001*** |
| PBS | -163472 | 11864 | -13.78 | < 0.001*** |
| Sham | -144213 | 11864 | -12.16 | < 0.001*** |
| Oenocyte | -365954 | 10772 | -33.97 | < 0.001*** |
| Plasmatocyte | -248740 | 10772 | -23.09 | < 0.001*** |
| Prohemocyte | -194008 | 10772 | -18.01 | < 0.001*** |
| GLMM model output / Hemocyte of 5-day-old bees at 22°C |  |  |  |  |
| (Intercept) | 327299 | 4858 | 67.37 | < 0.001*** |
| CLD | -109201 | 5432 | -20.10 | < 0.001*** |
| LP | -106597 | 5432 | -19.62 | < 0.001*** |
| PBS | -76042 | 5432 | -14.00 | < 0.001*** |
| Sham | -61319 | 5432 | -11.29 | < 0.001*** |
| Oenocyte | -216417 | 4858 | -44.55 | < 0.001*** |
| Plasmatocyte | -138556 | 4858 | -28.52 | < 0.001*** |
| Prohemocyte | -70889 | 4858 | -14.59 | < 0.001*** |

**Table 3.** Generalized Linear Model (GLMM) with mixed effects, predicting the effects of treatment and bacterial challenge on total hemocyte count. The model was conducted on hemocyte per mL data as a response variable and both treatment and hemocyte cell type as predictor variables: (GLMM <-response variable ∼ Treatment, data = x, family = quasi), for 15-day-old bees at 22°C. Levels of significance are *p* □< □ 0.001***.

| GLMM model output / Hemocyte of 15-day-old bees at 22°C |  |  |  |  |
| --- | --- | --- | --- | --- |
| 22°C | Estimate | Std Error | T-Value | P-Value |
| (Intercept) | 186285 | 5793.3 | 32.15 | < 0.001*** |
| CLD | -86424 | 8193 | -10.54 | < 0.001*** |
| CLD_E.coli | -116424 | 8193 | -14.21 | < 0.001*** |
| CLD_S.aureus | -114132 | 8193 | -13.93 | < 0.001*** |
| Control_E.coli | -47083 | 8193 | -5.74 | < 0.001*** |
| Control_S.aureus | -50312 | 8193 | -6.14 | < 0.001*** |
| LP | -79201 | 8193 | -9.66 | < 0.001*** |
| LP_E.coli | -110625 | 8193 | -13.50 | < 0.001*** |
| LP_S.aureus | -110208 | 8193 | -13.45 | < 0.001*** |
| PBS | -63160 | 8193 | -7.70 | < 0.001*** |
| PBS_E.coli | -95694 | 8193 | -11.68 | < 0.001*** |
| PBS_S.aureus | -95729 | 8193 | -11.68 | < 0.001*** |
| Sham | -55556 | 8193 | -6.78 | < 0.001*** |
| Sham_E.coli | -86389 | 8193 | -10.54 | < 0.001*** |
| Sham_S.aureus | -94583 | 8193 | -11.54 | < 0.001*** |

## Acknowledgment

Author contributions

Conceptualization: MO, DK, SK, and MA; Data Curation: MO, DK and SK; Formal analysis: MO, MA and SK; Funding acquisition: SK; Investigation: MO and SK; Methodology: MO and SK; Project Administration: SK; Resources: SK, MG, JA; Supervision: SK; Writing -Original Draft: MO and SK; Writing-Review & editing: MO, SK, JA, MG, MA. All authors have read and approved the content of this manuscript.

