## Supplemental Figure S1-S4; Table S1 for "Clodronate liposomes untangle the role of hemocytes in *Apis mellifera* response to temperature variation and microbial infection"

### Supplementary Materials

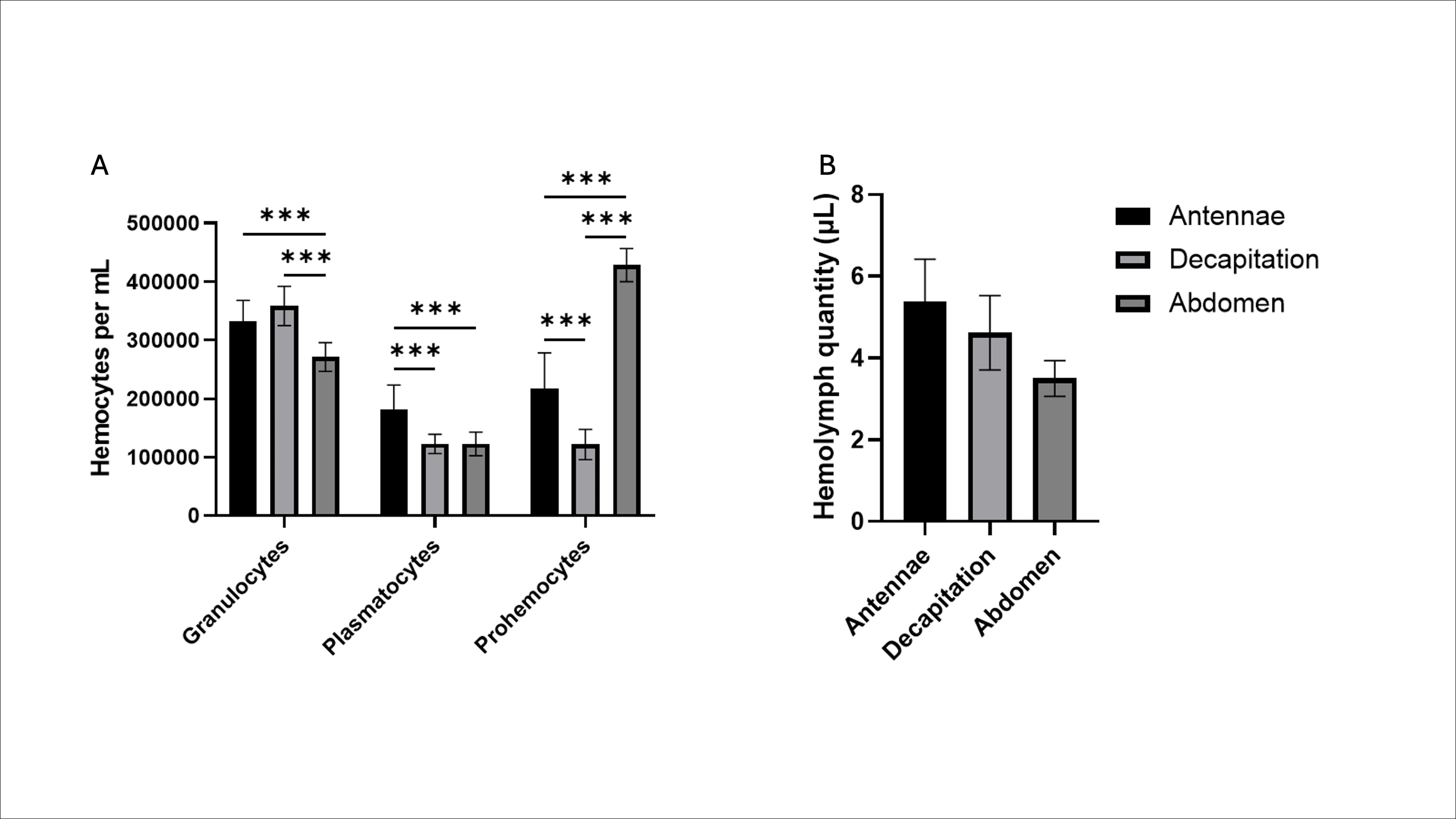

**Figure S1. Comparison of Hemolymph Extraction Methods.** Differential hemocyte type abundance (A) and hemolymph quantity in µL (B) in the varying hemolymph extraction of individual bees. Levels of significance: *p* < 0.05*, *p* < 0.01**, *p* < 0.001***.

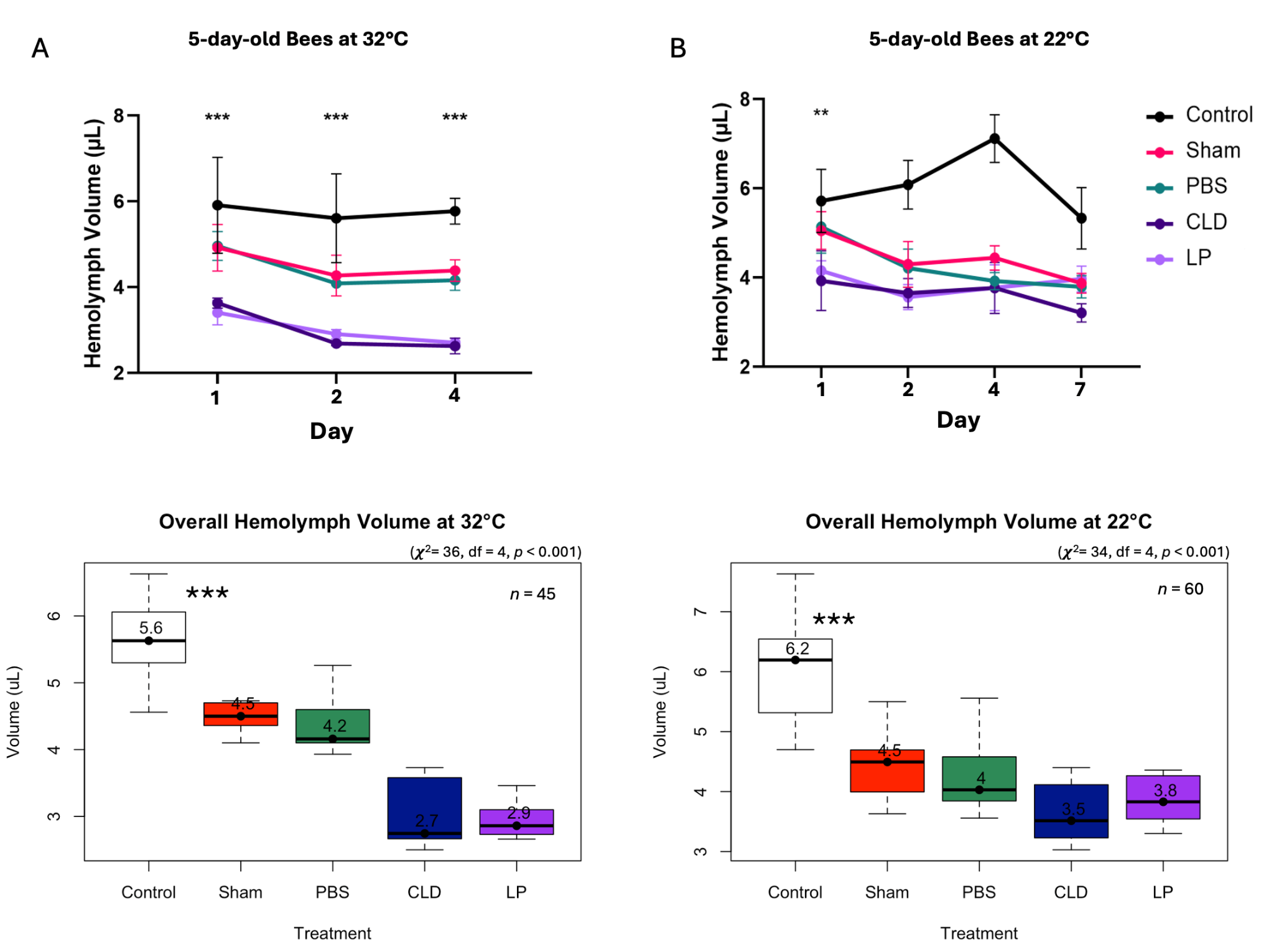

**Figure S2. Effect of injection treatment on hemolymph volume in 5-day-old bees at 32°C and 22°C.** Over time, hemolymph volume (μL) yield for bees housed at 32°C (A) and 22°C (B). Statistical differences between groups were determined using the Kruskal-Wallis test. Median is displayed for each boxplot and asterisks mark the levels of significance (*p* < 0.05*, *p* < 0.01**, *p* < 0.001***). Error bars in line graphs represent the standard deviation (SD).

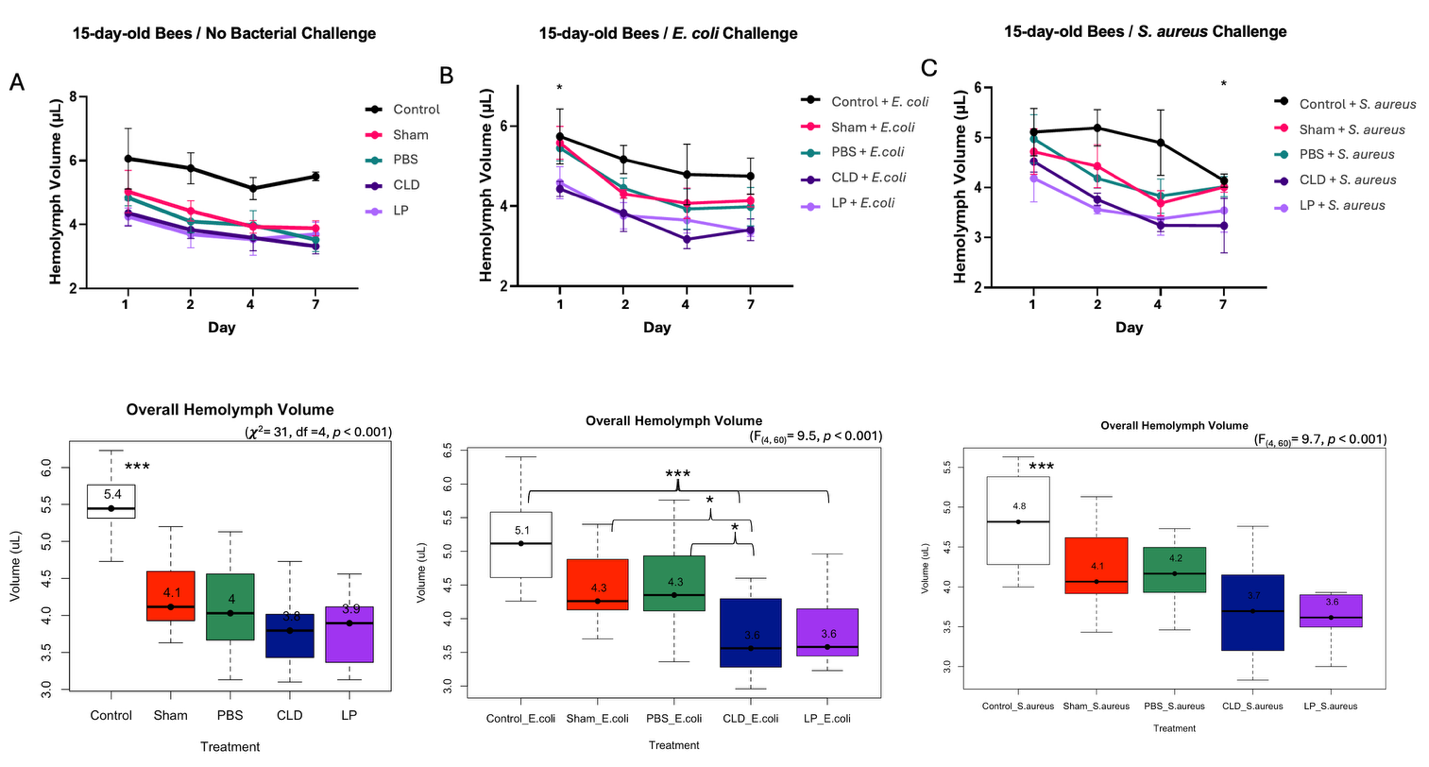

**Figure S3. Effect of injection treatment on hemolymph volume in 15-day-old bees.** Hemolymph volume (μL) yield over time for bees housed at 22°C with no bacterial challenge (A), (B) challenged with *E. coli*, and (C) with *S. aureus*. Statistical differences between groups were determined using one-way ANOVA and Kruskal-Wallis. Median is displayed for each boxplot, and asterisks mark the levels of significance (*p* < 0.05*, *p* < 0.01**, *p* < 0.001***). Error bars in line graphs represent the standard deviation (SD).

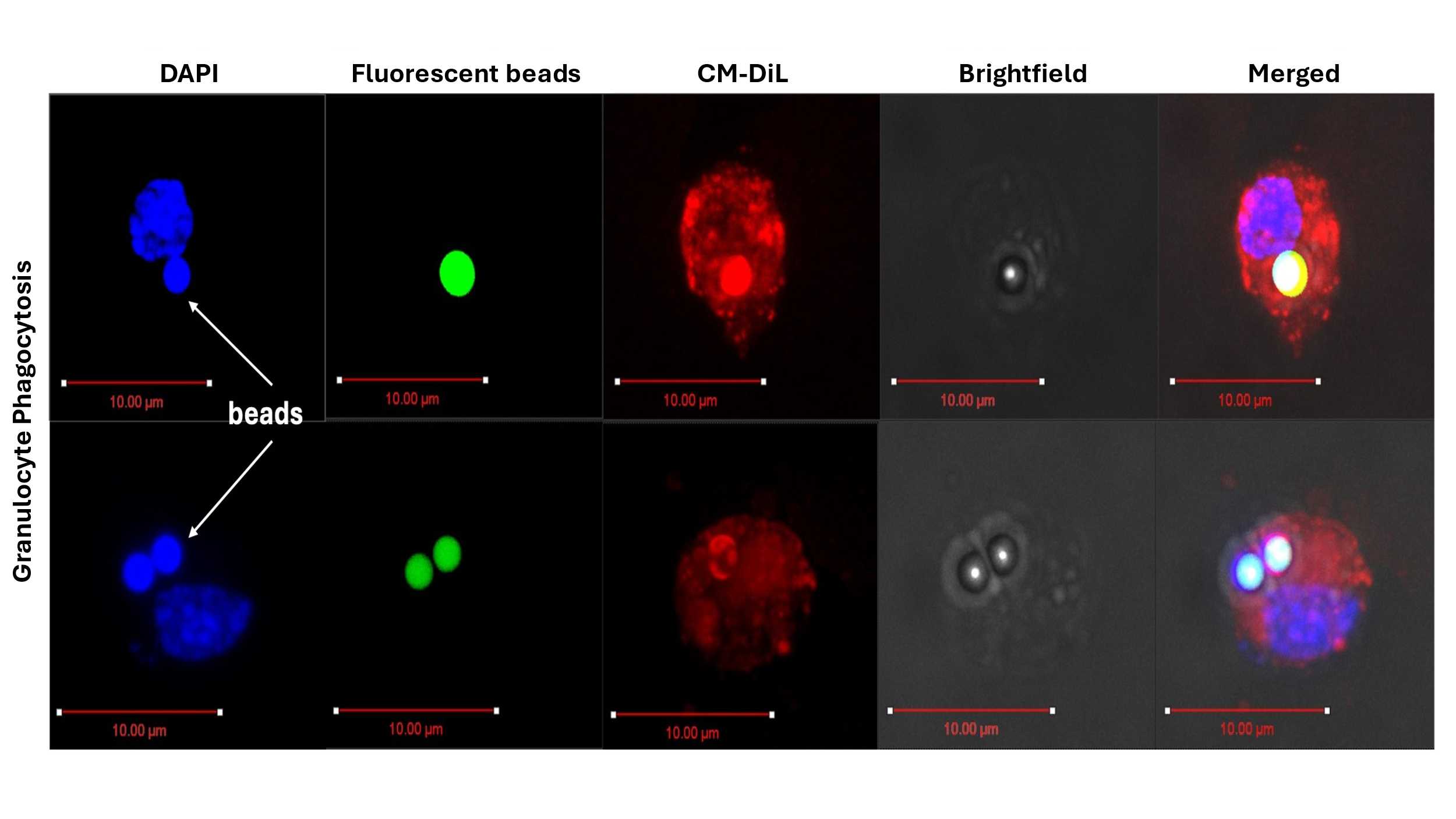

**Figure S4: Visualization of phagocytosis in adult honey bee granulocytes by fluorescent confocal microscopy.** Honey bees were anesthetized and injected with green fluorescent carboxylate beads and 0.75mM Vybrant CM-DiL lipophillic stain and then incubated for 30 minutes at 32°C to allow for phagocytosis of the beads and absorption of CM-DiL. Hemolymph was then extracted, and hemocytes were fixed, permeabilized, and stained with DAPI. Phagocytosis of the fluorescent beads allowed for confirmation that the CLD could also be phagocytized by individual granulocytes. The extreme brightness of the beads saturated the detector at minimum exposure settings, inducing crosstalk between fluorescent channels.. Fluorescent beads 2µm in diameter were engulfed by granulocytes. Scale bar = 10µm.

**Table S1.** **Summary of hemocyte quantity per treatment and bacterial challenge in 15-day-old bees.**

|  | **15-day-old bees at 22C** | | | | |
| --- | --- | --- | --- | --- | --- |
| **Granulocyte** | **Day 1** | **Day 2** | **Day 4** | **Day 7** | **Average** |
| Control | 331666 | 333888 | 321666 | 298333 | 321388 |
| Control + E.coli | 307222 | 233333 | 220000 | 161666 | 230555 |
| Control + S.aureus | 307222 | 232222 | 212222 | 166666 | 229583 |
| Sham | 254444 | 212222 | 228888 | 217777 | 228333 |
| Sham + E.coli | 237777 | 153333 | 150555 | 145000 | 171666 |
| Sham + S.aureus | 217777 | 121666 | 155555 | 131111 | 156527 |
| PBS | 233888 | 185000 | 201111 | 195000 | 203750 |
| PBS + E.coli | 213888 | 130000 | 151666 | 140555 | 159027 |
| PBS + S.aureus | 211666 | 133333 | 160555 | 123333 | 157222 |
| CLD | 205000 | 150555 | 167777 | 143333 | 166666 |
| CLD + E.coli | 161111 | 105000 | 132222 | 105000 | 125833 |
| CLD + S.aureus | 147222 | 118888 | 142222 | 111111 | 129861 |
| LP | 213333 | 170000 | 168888 | 149444 | 175416 |
| LP + E.coli | 155000 | 132777 | 128333 | 116111 | 133055 |
| LP + S.aureus | 149444 | 130000 | 120555 | 123888 | 130972 |
| **Plasmatocyte** |  |  |  |  |  |
| Control | 185000 | 186111 | 185555 | 155555 | 178055 |
| Control + E.coli | 182222 | 154444 | 123888 | 75000 | 133888 |
| Control + S.aureus | 180555 | 143333 | 127222 | 72777 | 130972 |
| Sham | 162222 | 118333 | 107777 | 107222 | 123888 |
| Sham + E.coli | 147777 | 95555 | 89444 | 65000 | 99444 |
| Sham + S.aureus | 137777 | 70555 | 85000 | 69444 | 90694 |
| PBS | 156666 | 125555 | 122222 | 107222 | 127916 |
| PBS + E.coli | 122777 | 70555 | 86111 | 62777 | 85555 |
| PBS + S.aureus | 117777 | 71111 | 92222 | 61111 | 85555 |
| CLD | 108333 | 108888 | 98333 | 91666 | 101805 |
| CLD + E.coli | 95555 | 62777 | 52222 | 46666 | 64305 |
| CLD + S.aureus | 82222 | 66111 | 53888 | 53888 | 64027 |
| LP | 126666 | 119444 | 102222 | 101666 | 112500 |
| LP + E.coli | 88333 | 79444 | 61111 | 72222 | 75277 |
| LP + S.aureus | 82777 | 72222 | 51666 | 55555 | 65555 |
| **Prohemocyte** |  |  |  |  |  |
| Control | 252222 | 200555 | 230555 | 180555 | 215972 |
| Control + E.coli | 225555 | 176666 | 151666 | 113333 | 166805 |
| Control + S.aureus | 231111 | 166666 | 151111 | 102777 | 162916 |
| Sham | 165000 | 144444 | 149444 | 156111 | 153750 |
| Sham + E.coli | 185555 | 97222 | 102777 | 68888 | 113611 |
| Sham + S.aureus | 148888 | 78333 | 100000 | 77222 | 101111 |
| PBS | 178888 | 136111 | 133333 | 121111 | 142361 |
| PBS + E.coli | 145000 | 82222 | 107222 | 82777 | 104305 |
| PBS + S.aureus | 144444 | 86666 | 107222 | 80000 | 104583 |
| CLD | 130000 | 112222 | 121666 | 106666 | 117638 |
| CLD + E.coli | 101111 | 73333 | 79444 | 60555 | 78611 |
| CLD + S.aureus | 86111 | 82777 | 91666 | 61666 | 80555 |
| LP | 159444 | 119444 | 124444 | 113333 | 129166 |
| LP + E.coli | 102777 | 67777 | 72222 | 91111 | 83472 |
| LP + S.aureus | 102777 | 91111 | 91666 | 86666 | 93055 |
| **Oenocytes** |  |  |  |  |  |
| Control | 23888 | 26111 | 42222 | 26666 | 29722 |
| Control + E.coli | 33888 | 32777 | 17777 | 17777 | 25555 |
| Control + S.aureus | 23888 | 26111 | 18333 | 13333 | 20416 |
| Sham | 13888 | 18888 | 18333 | 16666 | 16944 |
| Sham + E.coli | 25555 | 13888 | 13333 | 6666 | 14861 |
| Sham + S.aureus | 34444 | 13888 | 19444 | 6111 | 18472 |
| PBS | 21111 | 27222 | 10555 | 15000 | 18472 |
| PBS + E.coli | 17222 | 12777 | 15555 | 8333 | 13472 |
| PBS + S.aureus | 22222 | 11111 | 18333 | 7777 | 14861 |
| CLD | 19444 | 11666 | 12777 | 9444 | 13333 |
| CLD + E.coli | 15555 | 5000 | 17777 | 4444 | 10694 |
| CLD + S.aureus | 14444 | 16666 | 18333 | 7222 | 14166 |
| LP | 12222 | 8888 | 12222 | 11666 | 11250 |
| LP + E.coli | 16111 | 12222 | 7777 | 7222 | 10833 |
| LP + S.aureus | 25000 | 13888 | 11666 | 8333 | 14722 |
